# Community-aware explanations in knowledge graphs with XP-GNN

**DOI:** 10.1101/2024.01.21.576302

**Authors:** Andrés Martínez Mora, Dimitris Polychronopoulos, Michaël Ughetto, Sebastian Nilsson

**Affiliations:** AstraZeneca, Gothenburg, Sweden; AstraZeneca & Ochre Bio Ltd, Cambridge & Oxford, United Kingdom

## Abstract

Machine learning applications for the drug discovery pipeline have exponentially increased in the last few years. An example of these applications is the biological Knowledge Graph. These graphs represent biological entities and the relations between them based on existing knowledge. Graph machine learning models such as Graph Neural Networks can be applied on top of knowledge graphs to support the development of novel therapeutics. Nevertheless, Graph Neural Networks present an improved performance at the expense of complexity, becoming difficult to explain their decisions. State-of-the-art explanation algorithms for Graph Neural Networks focus on determining the most relevant subgraphs involved in their decision-making while considering graph elements (nodes and edges) as independent entities and ignoring any communities these graphs could present. We explore in this work the idea that graph community structure in biological Knowledge Graphs could provide a better grasp of the decision-making of Graph Neural Networks. For that purpose, we introduce *XP-GNN*, a novel explanation technique for Graph Neural Networks in Knowledge Graphs. XP-GNN exploits the communities of nodes or edges in graphs to refine their explanations, inspired by *cooperative game theory*. We characterize XP-GNN in a basic example and in terms of scalability and stability. In two relevant use cases for the drug discovery pipeline, XP-GNN provides more relevant explanations than previous techniques, being evaluated quantitatively and by domain experts. At the same time, XP-GNN presents limitations on scalability and stability, which we will address.

**ACM Reference Format:** Andrés Martínez Mora, Dimitris Polychronopoulos, Michaël Ughetto, and Sebastian Nilsson. 2024. Community-aware explanations in knowledge graphs with XP-GNN. In *Proceedings of ACM Conference (Conference’17)*. ACM, New York, NY, USA, 21 pages. https://doi.org/10.1145/nnnnnnn.nnnnnnn

**This work has been funded by AstraZeneca AB, Mölndal, Sweden and AstraZeneca Cambridge. Unfortunately, due to proprietary reasons from AstraZeneca AB, the data used in this work cannot be shared**.

## 1 INTRODUCTION

Launching novel therapies in the pharmaceutical industry has an average cost of two billion US dollars and takes in average fourteen years [1]. Many machine learning frameworks have been designed to accelerate and reduce the cost of novel therapies, thanks to the increasing quantities of data generated every day in the drug discovery pipeline. An example of ML technique automating the generation of new knowledge is the biomedical *knowledge graph* (KG). Biomedical KGs represent biological entities (genes, diseases, compounds, etc.) as nodes and their relations between them as edges connecting these nodes. KGs can be analyzed with ML models such as Graph Neural Networks (GNNs) to generate novel insights.

The drug discovery pipeline is tightly regulated to ensure that novel therapies are safe and effective. Consequently, any process in a ML model should be explainable, providing both the model decision and the background of this decision. Explainable models may bring improvements in user experience, continuous model enhancement, and root-cause analysis of their defects. GNNs in biological KGs suffer from lack of transparency, with many efforts recently focused on combining GNN predictability with explainability [2–6]. These works focus on the extraction of the most relevant nodes and edges that approximate the behavior of the GNN to the complete graph, treating these nodes and edges in isolation.

Nevertheless, the represented entities in biological KGs conduct shared functions and rarely work in isolation, as algorithms such as [2–6] model. For example, the represented genes in a biological KG work toward a common goal in communities known as *pathways*. We want to study whether these communities would better encode the decision-making process of GNNs in these graphs rather than using isolated graph elements in the explanation process. For that purpose, we present *XP-GNN*, a technique that aims to provide more relevant explanations for GNNs in biological KGs by benefitting from graph communities, inspired by *cooperative game theory*.

### Our contributions

1. We propose an explanation setup for GNNs inspired by cooper-ative game theory.
2. We design XP-GNN, an algorithm that efficiently approximates node or edge explanations from prior information from graph communities.
3. We characterize XP-GNN in simple examples in terms of scalability and stability.
4. We execute and evaluate XP-GNN quantitatively and with the help of biology experts, in two real-world use cases in a biological KG.

The remaining of this paper has the following structure. Section 2 contains an overview of related work for biological KGs, graph neural networks, and explanation resources for machine learning. Section 3 includes notations and formal definitions for the basis of our setup. The XP-GNN framework design is introduced in Section 4. Empirical properties of XP-GNN attributions are discussed in Section 5. Section 6 describes the application of our setup in two drug discovery cases. The paper concludes with Section 7, where our work is summarized, and we enumerate limitations to be studied in future work.

## 2 RELATED WORK

### 2.1 Biological Knowledge Graphs

Biological Knowledge Graphs (KGs) represent available knowledge in biology as a graph with nodes and edges to generate new knowledge for the introduction of novel therapies. Biological KGs contain nodes representing biological entities (genes, diseases, compounds, among others) and edges that relate these concepts. These edges are based on the current knowledge in Biology. The basic unit of a KG is the *triple*, where a source concept *(s)* is linked to a target concept *(t)* by a relation edge *(r)*. KGs are most often *heterogeneous graphs*, as they tend to contain diverse types of nodes (*concepts*) and edges (*relations between concepts*).

Biological KGs are built with information from different sources, as can be various biomedical databases or scientific articles mined through *Natural Language Processing*. These sources have to be complemented to construct the most possible complete graph. This also implies a high need for standardization towards a common framework, which may introduce noise in the knowledge mining process. Moreover, any noise in the original data sources used to build the graph may propagate to the graph. Biological KGs can reach large sizes (i.e. thousands of nodes and millions of edges), requiring highly scalable models and limiting the application of techniques that could entail a high computational cost. Apart from these problems, KGs follow the *open-world theory*, where inexistent edges between two nodes do not necessarily imply that those nodes are unrelated, but rather that the relations are still unknown. Consequently, the models that process a KG always deal with a certain degree of uncertainty. The recent literature contains a range of biological knowledge graphs for novel therapy discovery while trying to alleviate the limitations presented in the previous paragraph [7–14].

### 2.2 Graph Neural Networks

Graph neural networks (GNNs) are machine learning algorithms that extract relevant information from graphs such as KGs to derive novel insights. GNNs are mathematical functions that learn embeddings or representations for nodes and edges in a graph so that close graph elements receive similar embeddings. GNNs are based on the assumption of *homophily*, where neighboring graph elements tend to fulfill a similar purpose and should be processed similarly. Homophily is mathematically constructed with a *message-passing* scheme, where node and edge embeddings are propagated between neighboring elements in the graph. This technique allows the propagation of graph information based on local graph structure. In each message-passing iteration, the embeddings from the different neighbors are processed by a feed-forward neural network and subsequently aggregated with some mathematical function (mean, sum, or other). The aggregated results are then mapped by a non-linear activation function such as ReLU to allow learning non-linearities [15–17]. Graph embeddings in GNNs can be randomly initialized, or from node or edge features that are already known. GNNs can learn about specific nodes (*node prediction*), specific edges (*link prediction*), or the given graph as a whole (*graph prediction*). Depending on the task, the learned embeddings from the message-passing layers are inputted to different functions that learn about nodes, edges, or graphs. In supervised learning, the predictions of the GNN are compared to the ground-truth labels by a loss function. The loss function is subsequently applied to update the model’s parameters through gradient descent. As with any machine learning model, GNNs depend on hyperparameters (number of layers, layer size, aggregation function, etc.) that are tuned in a validation set.

Many improvements have been suggested from the basic GNN implementation to improve model performance. Graph Attention Networks (GAT) consider that different neighbors in the message-passing phases may influence differently, and hence aim to learn the weights of these neighbors [18]. Graph Isomorphism Networks (GIN) learn the aggregation functions from the neighbors in the message-passing phase, with the aim to distinguish subgraphs with close substructures [19]. APPNP (Approximated Personalized Propagation of Neural Predictions) [20] shifts from a pure message-passing paradigm to a paradigm based on initially obtaining the predictions (without any neighbor information sharing) and then propagating these predictions across the graph based on *Personalized Page Rank*, a technique that estimates the relevance of each node in the graph [21]. Other improvements deal with the treatment of heterogeneous information, where the different types of nodes and edges are treated differently in the message-passing iterations.

### 2.3 Explainability in Machine Learning

Understanding the decision-making process of machine learning models such as GNNs is critical, especially in drug discovery, as any action taken requires proper justification. Machine learning predictions alone are insufficient, so explainability becomes crucial. Explainable models provide also a better grasp of how they process the input data, improving user experience and traceability, since the user can better understand why some models could fail.

Linear models and decision trees are inherently explainable machine learning models, since the parameters governing these models tend to be directly related to the way the models process the input data. More complex models sacrifice explainability in exchange for performance, requiring *explanation algorithms* that can be applied on top of them to help understand how they make decisions. There are different categories of explanation models:

- *Global vs. local algorithms*. Global algorithms aim to explain models for any data instance. It is common to assess linear or tree-based models with these techniques, where the global importance of each feature is computed [22]. Local algorithms work only with individual data instances and are usually simpler to compute. GNNs are usually analyzed with local methods on a set of samples of interest, as they are computationally less expensive.
- *Model-specific vs. model agnostic algorithms*. Model-specific algorithms are designed to explain specific architectures. These techniques often rely on the gradients used to update the model during training. This is the case for class-activation maps or integrated gradients for Convolutional Neural Networks [23], for example. Model-agnostic algorithms work with any architecture and are preferred for GNNs, as they are independent of any mechanisms that the GNNs use in manipulating the models. A popular model-agnostic technique is LIME [24], where an interpretable linear model is fit in the vicinity of a data instance from the explained model.
- *Rule-based algorithms*. These algorithms provide explanations based on causality relations. They are rarely implemented, as these relations are usually unavailable. These methods can be useful for GNNs, through complementary techniques such as reinforcement learning [25].

#### 2.3.1 Co-operative Game Theory Based Methods

Explainability techniques can be also built from *Game Theory*. Game Theory methods are model-agnostic, representing the data elements to be explained as *players* competing in a *game* to obtain a *reward*. This reward will be the output from the explained model, while the players can be data features, or nodes or edges in a graph. With these tools, it is possible to construct a model explanation that informs of the most relevant features, nodes, or edges based on how these elements cooperate to obtain the model output.

One of the most applied tools in game theory for machine learning model explainability is the *Shapley value*, which computes the contribution of each player in the game under the assumption that all players are independent[26]. Shapley value theory computes the average change in reward when a player is considered in the game, compared to when that same player is absent. The Shapley value considers all player combinations where the player of interest is absent, comparing the model output with these combinations to the model output for the same combinations including the player of interest. Each of the considered combinations is known as *player coalition*. Fig. 2 reflects the computation of the Shapley value for a *star* in a player set *P* composed of three other shapes. These shapes interact with each other to obtain a reward $ by means of a reward function *f*. The Shapley value for the star considers all the coalitions of shapes where the star is present and compares evaluating *f* when the star is present as opposed to when the star is absent. The average of the differences in each evaluated coalition provides the Shapley value for the star, considering the number of non-star shapes in the coalitions excluding the stars, *S*.

**Figure 1.**
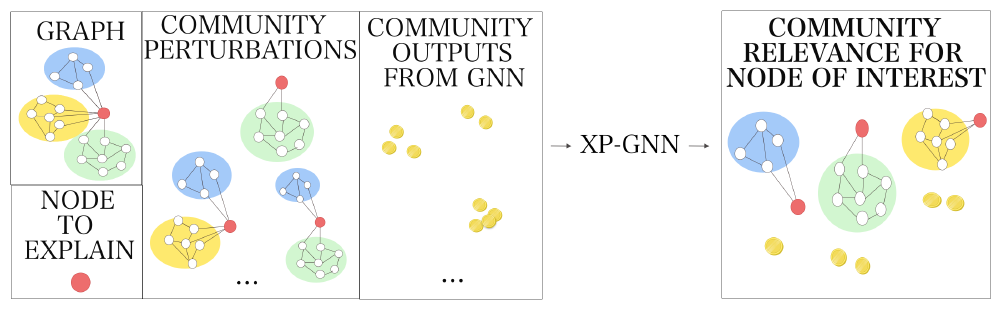
An input graph with colored communities is perturbed according to its community structure for the node to explain. The perturbations have a certain GNN output. The GNN outputs are used by *XP-GNN* to determine the actual contribution of each community in the GNN for the node of interest.

**Figure 2.**
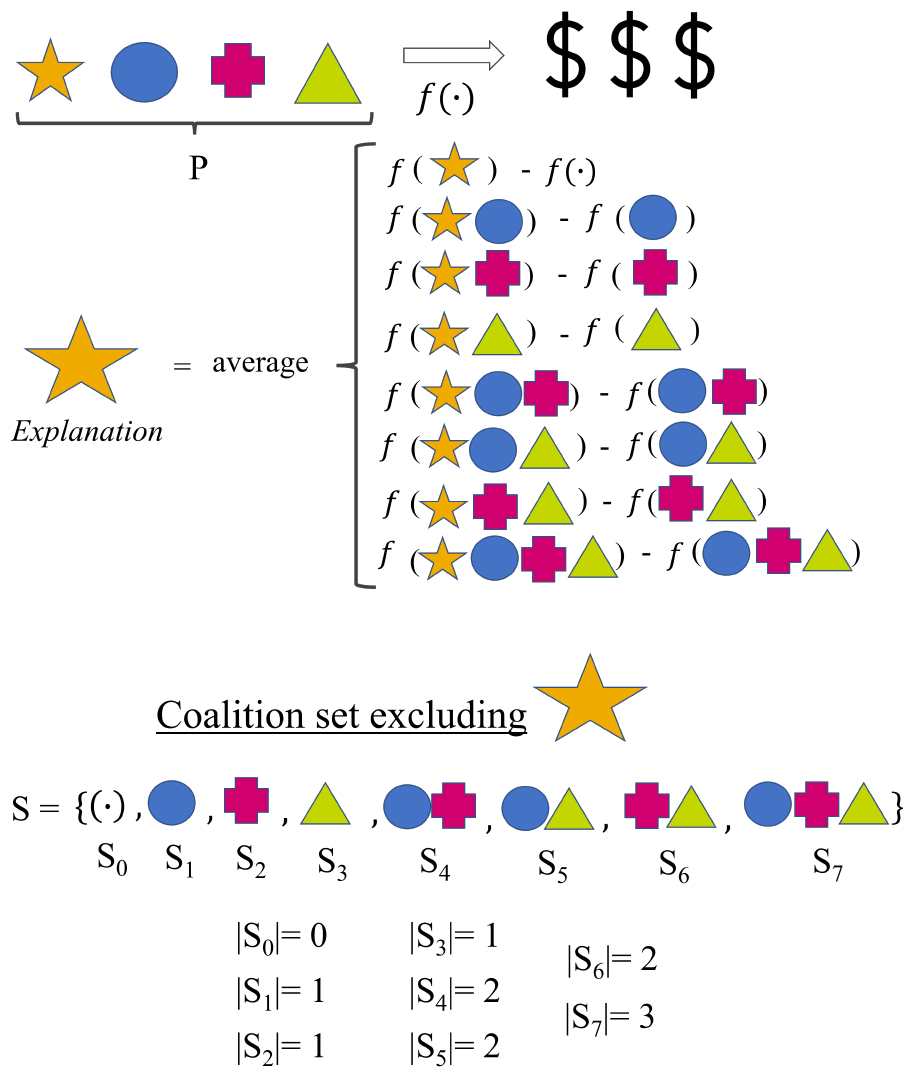
Considered coalitions to compute the Shapley value for the star shape. Shapley value theory considers four players *P* that interact with a model *f* to obtain a reward ($$$). We include the coalitions *S* from the complete permutation set Π, excluding the star shape. The number of player in each coalition *S* is |*S* |.

To explain machine learning models, the coalitions of players can be considered as combinations of features, nodes, or edges. In the case of feature coalitions, the absent features are masked to some reference value (a mean value, zero, or some other value). In the case of nodes or edges, the absent elements can be removed from the original graph, resulting in *graph perturbations*. The data points with masked features or the perturbed graphs are then inputted to the model to explain. Consequently, the model outputs are compared when that feature, node, or edge is present against to when the feature, node, or edge is masked or perturbed. Each computed difference is weighted according to the number of permuted elements and the total number of features, nodes, or edges considered.

When analyzing datasets with dozens or hundreds of features, edges, or nodes, the number of coalitions to analyze becomes very large. Consequently, the computation of the exact Shapley value is expensive, with many approximations suggested [27]. An extended acceleration method for the Shapley value is SHAP[28], where only a random set of all the potential coalitions is considered. SHAP constructs a mask with zeros and ones for those features, nodes, or edges that are either removed or considered. This mask is fit against the model outputs for the sampled coalitions through weighted linear regression. The coefficients of the regression model (also known as *surrogate model*) are the approximations of the Shapley value. For each sampled coalition, SHAP computes a kernel value as the weight for the surrogate model, which depends on the number of active features, nodes, or edges in each coalition and the total number of elements in the analysis.

Shapley value theory assumes player independence, which is not always the case in the real world. Sometimes, the features, nodes, or edges can form communities that interact with the model to explain. These communities can present features, nodes, or edges in several communities at the same time. Therefore, the coalitions must consider these structures to provide a community-aware explanation. The *configuration value* has been proposed to process community-aware games [29]. For each community, it considers the following types of coalitions:

- *Internal coalitions (* ℑ ). They are only constructed with the players of the same community. In Fig. 3, it is aimed to compute the configuration values for the star shape, so the first two coalitions are *internal*, as these coalitions consider only players from *Community 1*.

**Figure 3.**
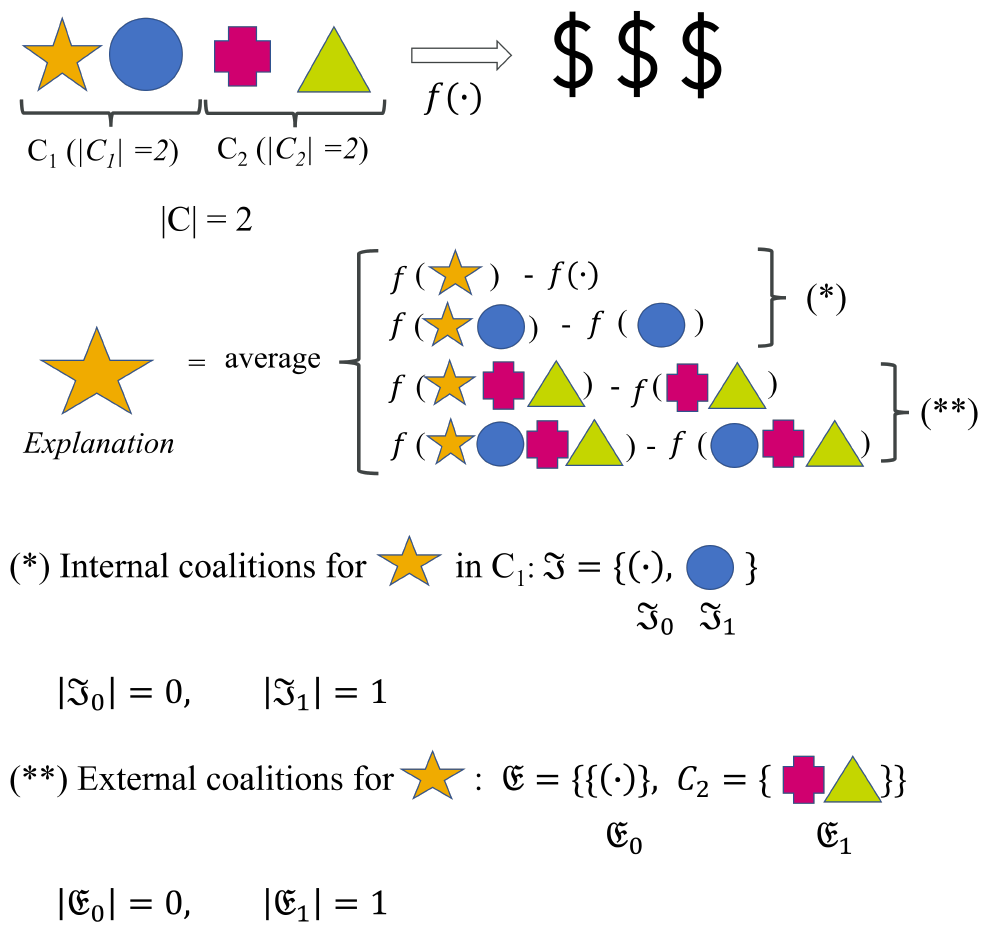
Considered coalitions to compute the configuration value for the star shape. Configuration value theory considers four players in a community set *C* that interact with a model *f* to obtain a reward ($$$). We include the internal coalitions ℑ and external coalitions 𝔈, with their respective sizes |ℑ| and |𝔈 |.
- *External coalitions (* 𝔈 ). These coalitions consider players from other communities apart from the community analyzed, known as *external communities*. If a player from an external community is introduced, all the other players in that external community should be considered, too. In Fig. 3, as the star shape is the studied subject, the analyzed community is *C*_1_. Consequently, external coalitions can be built with the cross or the triangle shapes in *C*_2_, but they always have to include *both* of them. As an important fact, the size of these coalitions equals the number of external communities considered, rather than the total number of players from these communities.

When many features, nodes, or edges with a community structure have to be analyzed, the configuration values can be also approximated with similar techniques as SHAP[28].

#### 2.3.2 Explanations of Graph Neural Networks

Several explanation algorithms are available for GNNs. They can explain nodes, edges, or complete graphs depending whether the problem is a node prediction, a link prediction, or a graph prediction problem, respectively. Most of the explanation approaches are local and focus on extracting the subgraph that is processed as similar as possible to the full graph by the GNN. Most often, the size of this subgraph is upperly bounded to avoid explanation subgraphs that are very similar to the original subgraph. The nodes and edges of the output subgraph are labeled with some score that indicates their relevance for the GNN output. In some cases, the scores are signed and can tell if the node or edge contributes for or against the behavior of the GNN.

[2] suggested *GNNExplainer*, an algorithm that maximizes the Mutual Information of the GNN behavior in a candidate subgraph against the complete graph for some node or edge. [3] aimed for more global explanations with *PGExplainer* by approximating the global edge distribution with a neural network and then sampling subgraphs from this distribution while optimizing for Mutual Information as GNNExplainer. [4] frames the graph explanation problem as a probabilistic problem, determining a first subgraph with chi-square statistics and then optimizing this subgraph with Probabilistic Graph Models in an approach named *PGMExplainer*. [5] explores with *SubgraphX* Monte Carlo Tree Search (MCTS) to construct candidate subgraphs that are refined through Shapley value estimation for graph nodes. Node-wise Shapley values are also computed as graph explanations in *GraphSVX* [6].

Many of these approaches have been applied in relatively small graphs, so it is uncertain how larger graphs as KGs could be processed, where many different types of nodes and edges can be found. Furthermore, these methods ignore the potential communities where the nodes or edges in the graph could be assembled, so that the provided explanations could be less linked to the structure of the graph itself.

## 3 PRELIMINARIES

### 3.1 Graph machine learning fundamentals

Graph machine learning tools extract insights from graph datasets such as KGs, for example, to find novel therapies. As presented in Section 2.1, KGs are composed of a node-set *V* with *v* nodes, an edge set ℰ with *e* edges connecting source and target nodes, a node type set (*T*), and an edge relation set (*R*).

#### Definition 1

***Input Knowledge Graph (KG)***

*G* = (*V*, ℰ, *T, R*, ***X***)

*with* 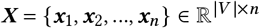 *a matrix with n node features. The number of features may vary for each node type*.

In graphs, node or edge players can arrange into overlapping communities. As mentioned in Section 1, these communities can provide prior information to build model explanations. A community set *C*, with *c* communities in a graph *G* can be defined as:

#### Definition 2

***Overlapping community set***.

*C* = {*C*_1_, …, *C* _*j*_, *C*_*m*_, …} *with* ◊(*C* _*j*_ ∩ *C*_*m*_ ≠ ∅) *and* 0 *< j, m* ≤*c, j* ≠ *m*

(♦ stands for *it is possible*).

The subset of communities where some node or edge *i* is located is an important piece in the computation of the configuration value of *i*, being defined as:

#### Definition 3

***Community subset for player*** *i*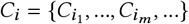*where C*_*i*_ ⊆ *C and* 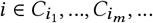

The remaining communities not containing node or edge *i* constitute the *external community subset*, defined as:

#### Definition 4

***External community subset for player*** *i*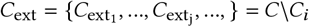*where C*_ext_ ⊆ *C, C*_ext_ ∩ *C* = ∅

To generate new knowledge from *G*, a GNN model *f*_θ_ applies a parameter set θ to extract relevant embeddings of nodes, edges, or graphs, as described in Section 2.2. *f*_θ_ is composed of several layers, where graph embeddings are propagated in each layer and then mapped by an activation function *σ*. In a layer *k*, the parameter set θ is composed by a weight set *W* ^(*k*)^ and a bias set *b* ^(*k*)^, while the embedding of a node *v* in a layer *k* can be encoded as 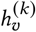. At the same time, the neighbors of *v* can be a set of nodes *u*, and their total number is *N* (*v*), also known as the *node degree* of *v*. With these concepts, the generic equation for a GNN is:

#### Definition 5

***Message-passing Graph Neural Network***, *k*^*th*^ ***layer***

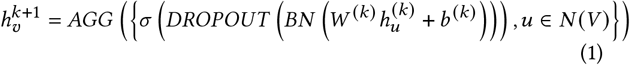

The first part of the equation reflects how the embeddings of the neighboring nodes *u* are aggregated, being regulated by *W* ^(*k*)^. The second part of the equation shows the influence of the embeddings in node *v* from the previous layer *k*, being regulated by *b* ^(*k*)^. This information can then be normalized through *Batch Normalization* (*∅N*) and passed through a *DROPOUT* operator. The aggregated information from all neighbors *u* and the information from previous layer *k* are non-linearly mapped by *σ*. A global aggregation operator (*AGG*) combines the non-linearly mapped information. *AGG* can be the sum, the maximum, or the mean. The equation above can be slightly modified for more complex GNN architectures [30].

### 3.2 Cooperative game theory

As introduced in Section 2.3, the game-theory concepts of *Shapley value* and *configuration value* can be used to construct explanations for GNNs as *f*, with the latter inputting information on graph structure. We can define a player set *P* :

#### Definition 6

***Player set***.

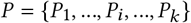

For GNN explanations, the players are the graph nodes or edges used to explain *f*. The number of players is |*P* |, with ∏ containing all their possible permutations. As introduced in Section 2.3, the Shapley value for player *i, ϕ*_*i*_, estimates the contribution of *i* as shown in Fig. 2 with the star shape. The model to explain is evaluated in player coalitions excluding this shape, comparing these outputs to the model outputs for the coalitions where the star is introduced. All the player coalitions excluding player *i* can be defined as Π \{*i*}. Any coalition or player subset in this set is *S, S* ⊆ Π \{*i*}. The number of players in the subset *S* is |*S* |. In the graph explanation paradigm, *S* would be a subset of nodes or edges sampled in a perturbed graph, to be later processed by a GNN. With all these elements, the Shapley value for a player *i* in a cooperative game is:

#### Definition 7

***Shapley value for player*** *i*

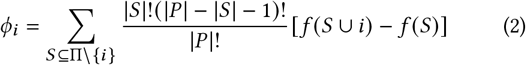

Shapley value theory comes with additional guarantees that motivate its preference for machine learning explanation algorithms:

#### Efficiency

the sum of the Shapley values for all players in *P* equals the value of the model *f* being explained. When a coalition that contains all players in *P* is considered, this is known as *grand coalition*

#### Symmetry

the Shapley value of a player *i* equals that of a player *j* if their marginal contributions (differences in value of *f* before and after considering them) is the same

#### Dummy player

If a player *i* does not contribute to any changes in the outputs of *f*, its Shapley value is zero.

#### Linearity

if linear relations exist between several players, their values can be linearly combined to obtain the resulting Shapley value.

The Shapley value considers all players in *P* as independent players (see Section 2.3). In case that players arrange in communities, the configuration value explains better how the model works given the communities the players form. Configuration value theory considers internal and external coalitions of players, based on the community set *C*, as explained in Section 2.3.

Internal coalitions involved in the computation of the configuration value include all player subsets in all communities in *C*_*i*_. In the graph explanation domain, *C*_*i*_ would be constituted by all graph communities containing node or edge *i*. Coalitions of internal players can be defined as:

##### Definition 8

***Internal coalition set***

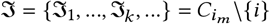

Where |ℑ_*k*_ | is the number of players considered in the *k*-th internal coalition (0 ≤ *k <* |ℑ|). In Fig. 3, ℑ contains the empty set and the circle shape. To explain graph-based models, ℑ_*k*_ would contain a perturbed graph with nodes or edges from the same community where node or edge *i* belongs.

External coalitions for the computation of configuration values are constructed with the players from *C*_ext_. In each coalition, the players of some of these communities are sampled, resulting in an *external coalition set* defined as:

##### Definition 9

***External coalition set***

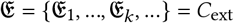

An external community that is sampled in 𝔈_*k*_ *must* introduce all the players in that community. In the example in Fig. 3, the players in *C*_2_ are considered in both external coalitions, as they are part of 𝔈_1_. In the graph explanation domain, 𝔈_*k*_ would contain a perturbed graph with nodes or edges from communities that do not include node or edge *i*, under the condition that all the nodes and edges from these selected communities are included.

Given all definitions above, the configuration value for a node or edge *i, ψ*_*i*_, given a community set *C* in a graph *G*, is defined as [29]:

##### Definition 10

***Configuration value for player*** *i*.

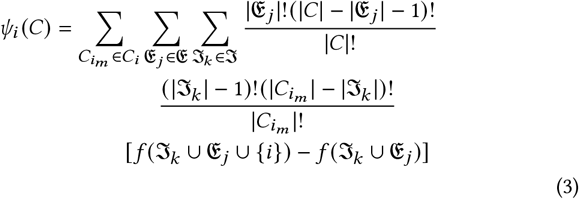

The equation iterates through all communities in the community subset of node or edge *i* (*C*_*i*_). For each community in *C*_*i*_, the algorithm iterates through the internal coalitions of nodes or edges, excluding *i* (ℑ). The equation then iterates through the coalitions of external communities in 𝔈. When evaluating the model to explain, the nodes or edges in the sampled coalitions are combined in ℑ_*k*_ ∪ 𝔈_*j*_.

## 4 THE FRAMEWORK DESIGN (METHODOLOGY)

In this section, we introduce the high-level design of XP-GNN, which was designed with the Python graph ML framework *PyTorch Geometric* [31]. All the source code for XP-GNN is available at: https://github.com/andres2631996/bikg_graph_explainability_public. XP-GNN receives the following inputs:

- A knowledge graph *G*
- An overlapping community set of graph elements (nodes or edges) *C*
- A GNN *y* = *f*_θ_ (*G*)
- An instance *s* to be explained. *s* can be a node *n* for node prediction, an edge *e* for edge prediction, or the graph *G* for graph prediction

KGs can have many nodes and edges, turning the direct computation of configuration values with Definition 10 as intractable. XP-GNN approximates *ψ* by sampling internal and external node or edge coalitions in *C* (as in the example in Fig. 3). These coalitions can be constructed with the perturbation of *G* into a set of perturbed graphs in *G*_perturbed_, to only contain the nodes and edges in the sampled coalitions. The graphs in *G*_perturbed_ are inputted to *f*_θ_ to obtain *y*_perturbed_ as outputs. A *kernel* score is determined for each perturbed graph, which depends on the perturbation degree from the original graph. These scores reflect the relevance of each graph perturbation to build the final explanation. As a last step, a local weighted linear regression model *g*_*ψ*_ is fit as a surrogate, inspired by [28]. The parameter set *ψ* is used to obtain the approximations of the configuration values. *g*_*ψ*_ is defined as:

### Definition 11

***Surrogate model for XP-GNN***

*g*_*ψ*_ : ℝ^*p*×|*V* |^ : ℝ^|*V* |^ *for node configuration values*

*g*_*ψ*_ : ℝ^*p*×|*E* |^ : ℝ^|*E* |^ *for edge configuration values*

Where *p* is the total number of graph perturbations. The outputs provided by XP-GNN are:

- The approximated configuration values (node-wise or edge-wise)
- The aggregated configuration values for the node or edge communities in *C*

The overall steps are represented in Fig. 4.

**Figure 4.**
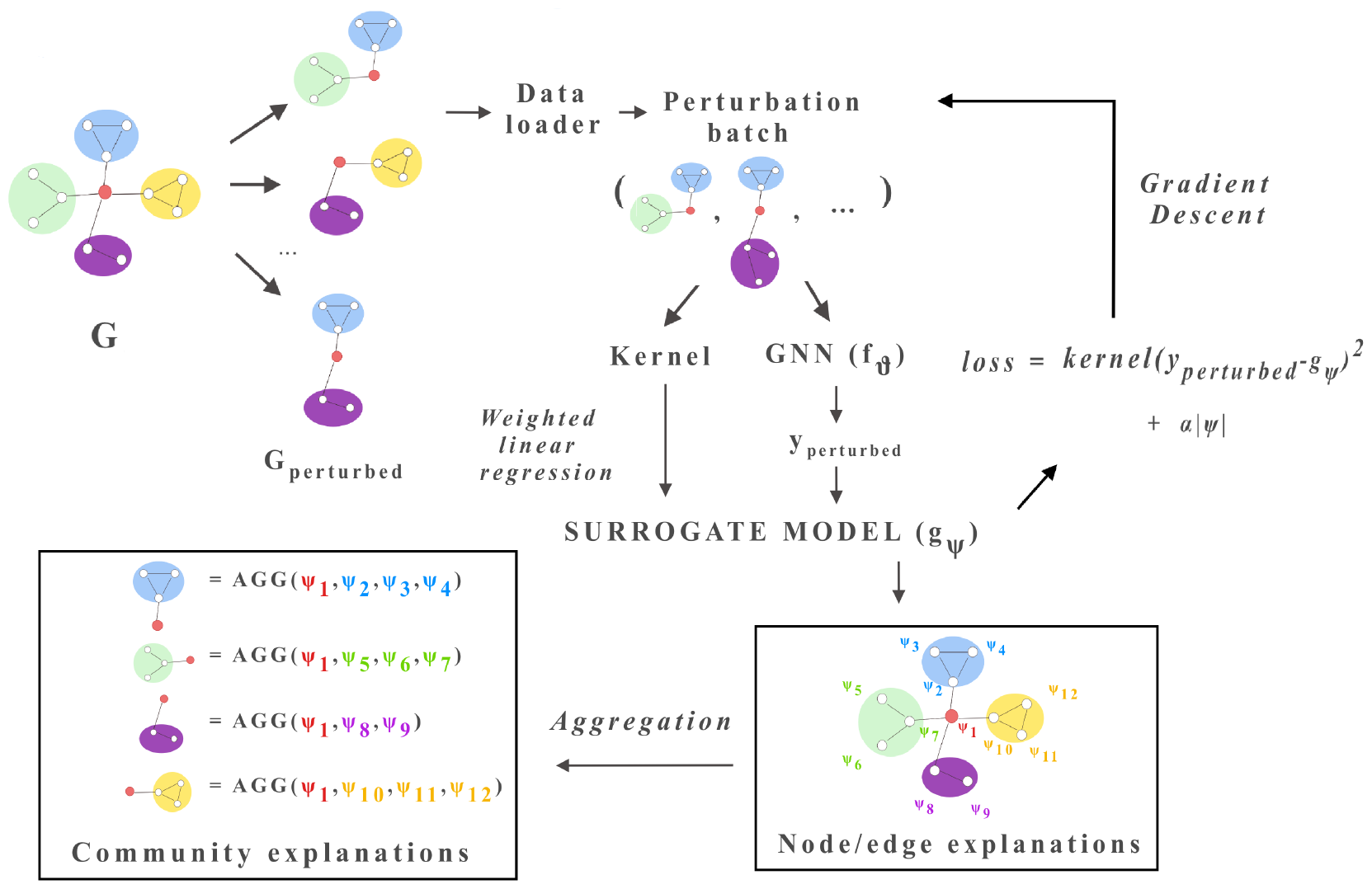
XP-GNN overview to explain the central node in *G*, with overlapping communities depicted in different colors. *G* is perturbed community-wise into *G*_perturbed_. This is stored in a dataloader, so that a surrogate model *g*_*ψ*_ can be trained iteratively, computing kernels and perturbed outputs for the perturbed graphs in the batch. *g*_*ψ*_ is a weighted linear regression model, optimized with Gradient Descent. *ψ* contains the node or edge explanations, which can be aggregated into community explanations.

### 4.1 Community-based graph perturbations

The input graph *G* is perturbed *p* times, so that *G*_perturbed_ contains *p* graphs with different perturbed nodes or edges. Those nodes or edges that do not make part of the sampled coalitions are removed. The sampled coalitions belong to the *internal* and *external* coalitions presented in Section 2.3. The complete graph perturbation process is as follows:

1. We iterate through all the communities in *C*.
2. In each community *c* ∈ *C*, we randomly sample its nodes or edges. These nodes and edges constitute the *internal coalitions*.
3. The remaining communities that are not considered in step 2 constitute the *external community set, C*_ext_. Some of these communities are sampled, constructing the *external coalitions* with *all* their respective nodes and edges.
4. The nodes and edges that are not sampled in any of the previous steps are removed, resulting in graph perturbations stored in *G*_perturbed_. The indexes of the preserved nodes or edges in each perturbed graph are applied to build a binary mask *M* ∈ ℝ^*p*×*n*^. *M* codes with ones the preserved nodes or edges and with zeros the removed graph elements. *n* is the total number of nodes or edges. If we build the explanations in terms of nodes, *n* is the number of nodes, while if we build the explanations with edges, *n* is the number of edges.

All graph perturbations are computed at once and stored in a data loader to train *g*_*ψ*_ in batch mode with gradient descent. If memory is limited, the coalition sampling process focuses on large communities only, as we hypothesize that most changes in the behavior of a GNN could be brought by large graph communities, so focusing only on those communities could save resources in resource-scarce situations. The approximation of the Configuration Value from Equation 3 is not violated in this situation, as all the possible coalitions of perturbed and unperturbed communities can never be sampled, and that happens even in cases where perturb all communities with a similar probability, and not only when focusing only on large communities, so there will always be slight deviations from the actual configuration values from Equation 3. This process allows to later sample few perturbed graphs to iteratively train a surrogate model that approximates locally the behavior of the GNN in the node, edge, or graph to be explained (see Section 4.4).

The node or edge indexes that are zeroed in mask *M* have to be removed for each perturbed graph in *G*_perturbed_. If 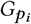 is a given perturbed graph, it is composed of a node feature matrix 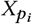 and an edge index matrix 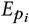.

If the columns in *M* represent nodes, we identify the nodes where *M* is zero and locate the edges where these nodes participate. These edges are removed from the edge index of *G*, resulting in 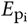. We can keep the same node feature matrix as the original graph as long as the edge indexes are perturbed.

If the columns in *M* represent edges, we identify the zeroed edge indexes, and directly remove them from the edge index of *G*, resulting 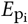. As happens with nodes, we can also keep the same node feature matrix as the original graph.

Perturbations are executed in batch mode (several at the same time), concatenating the node and edge sets of the graphs in *G*_perturbed_.

### 4.2 Forward Pass and Scoring

The number of sampled nodes or edges in each perturbation (*n*_permuted_) is applied to compute the *kernel* of that perturbation in the surrogate model (*K* ∈ ℝ^*p*^). *K* is computed as below, inspired by the *kernelSHAP* technique in [28]:

#### Definition 12

***Kernel values from kernelSHAP***

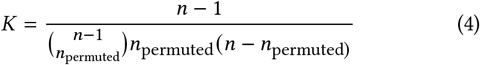

*n* is the number of nodes or edges in the original graph. The equation above is computed for several perturbations in batch mode. The perturbed graphs in *G*_perturbed_ are inputted to the GNN, obtaining:

#### Definition 13

***Output perturbations***

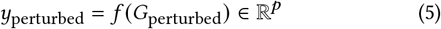

*yperturbed* contains node values for node prediction problems, edge values for edge prediction problems, and graph values for graph prediction problems. The predicted values in *yperturbed* are a continuous quantity, either actual predicted values in regression problems or predicted probabilities in classification problems, making *XP-GNN* an agnostic algorithm for regression or classification tasks.

### 4.3 Attribution estimation

The surrogate model *g*_*ψ*_ is iteratively fit with the perturbations from mask *M*, being optimized in gradient descent to learn *y*_perturbed_. XP-GNN has the number of the sampled perturbations (*m*) and the number of training iterations (*epochs*) as hyperparameters. The loss function to train *g* is a weighted mean square error loss with L1 regularization and regularization constant *α*, to obtain smoother distributions of the approximated configuration values:

#### Definition 14

***Weighted mean square loss***

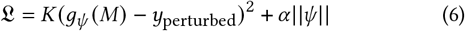

The learned coefficients of *g, ψ*, are the final approximations of the node-wise or edge-wise configuration values. These values can be aggregated into community-wise values *ψ*_comm_ with *ψ*_comm_ = *AGG* (ψ_*c*_). *AGG* is an aggregation operator commonly set to the mean or the median of the scores to avoid community-size bias (XP-GNN outputting large communities as the most relevant ones just because of their size).

### 4.4 Algorithm

#### Algorithm 1

Community-aware explanations for GNNs with XP-GNN

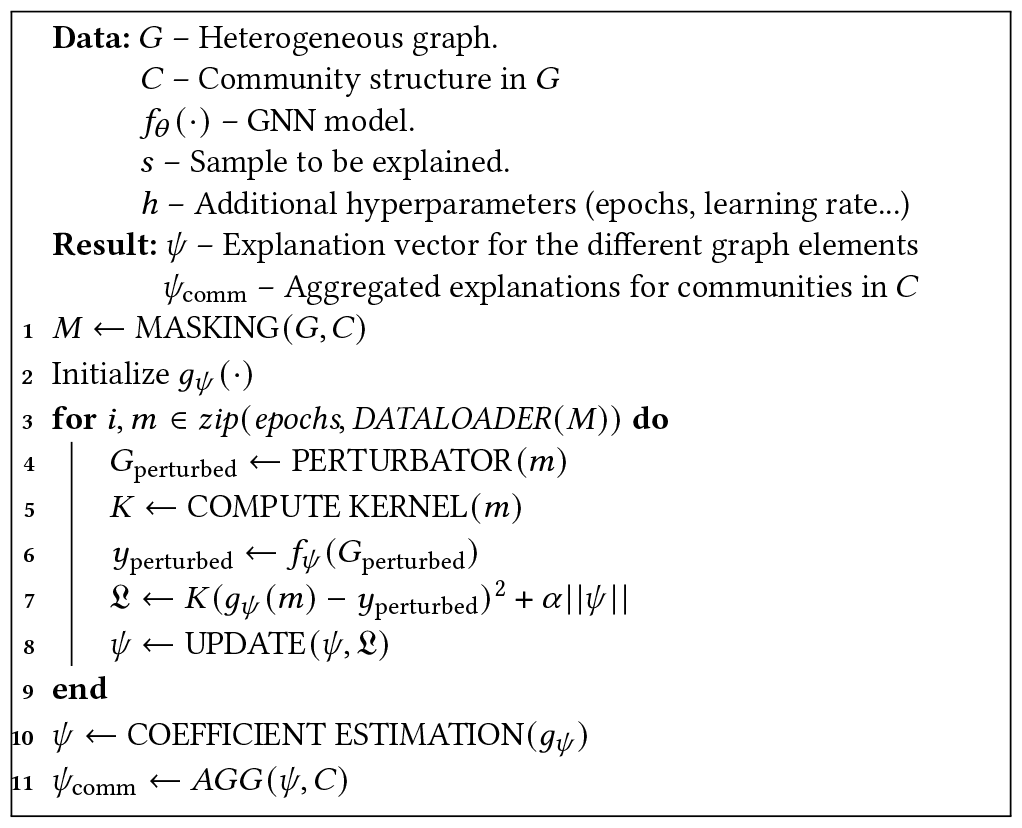

## 5 EXPERIMENTAL EVALUATION

This Section characterizes XP-GNN in settings with simple graph datasets and in the *Biological Insights Knowledge Graph* (*BIKG*) [32], a Biological KG owned by AstraZeneca. BIKG is comparable to PrimeKG, a popular biological KG that has been recently developed[14].

### 5.1 Synthetic example

We designed a simple example graph with five nodes, eight edges, random node features, and two node communities. One community contains just a node, while the other community contains three nodes. The graph has a central node outside these communities to observe how their behavior influences this central node. To examine the influences of the communities on the central node, two cases were simulated:

- *Case A*, where the three-node community (in blue in Fig. 7) is set to a label with value 0.1, while the central node (in black in Fig. 7) and the node in the small community (in red in Fig. 7) are set to 0.9 as label value, in a clear reference to the fact that the second community would present a high influence in the central node, and would be expected to have a high relevance in the explanation from *XP-GNN*.

**Figure 7.**
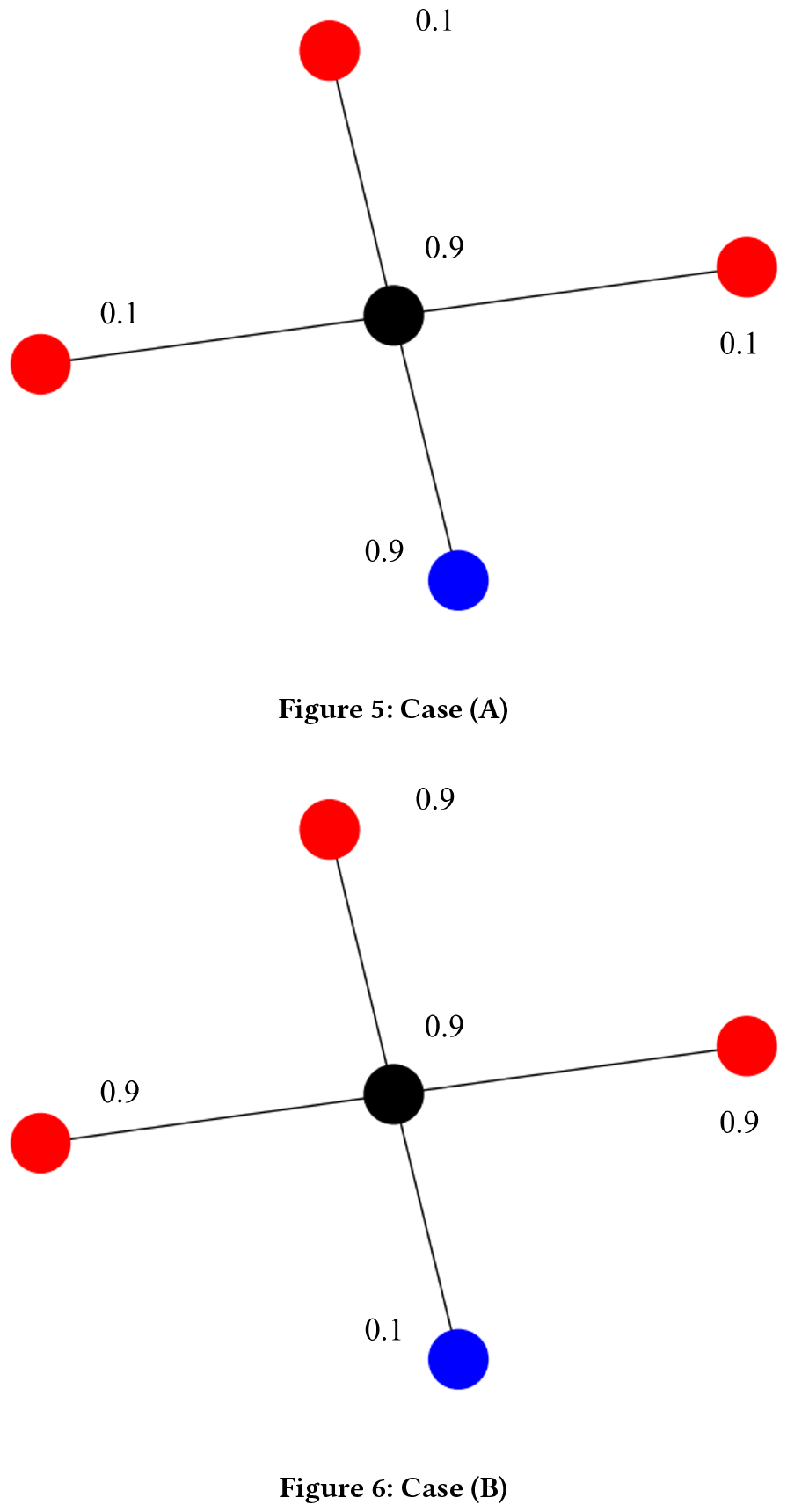
Cases A and B for toy example, with the same graph structure, but different node labels, to demonstrate the influence that the different communities (in red and blue) may exert in each case in the central node, in black.

**Figure 8.**
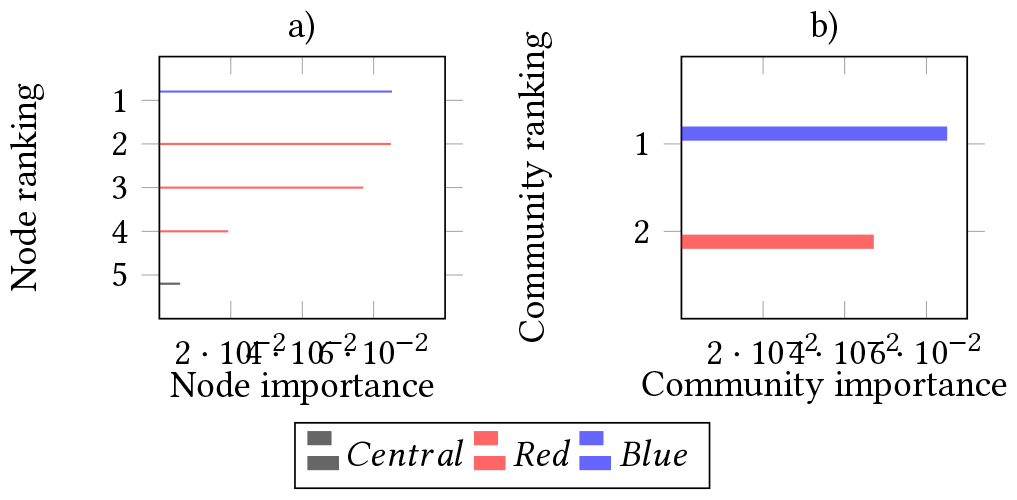
a) Node scores for the central node in the graph in Fig. 7a), when using a GNN to predict node-wise labels. Node bars receive the color of the community to which each node belongs. The central node is colored in black. b) Aggregated scores for the communities in Fig. 7.

**Figure 9.**
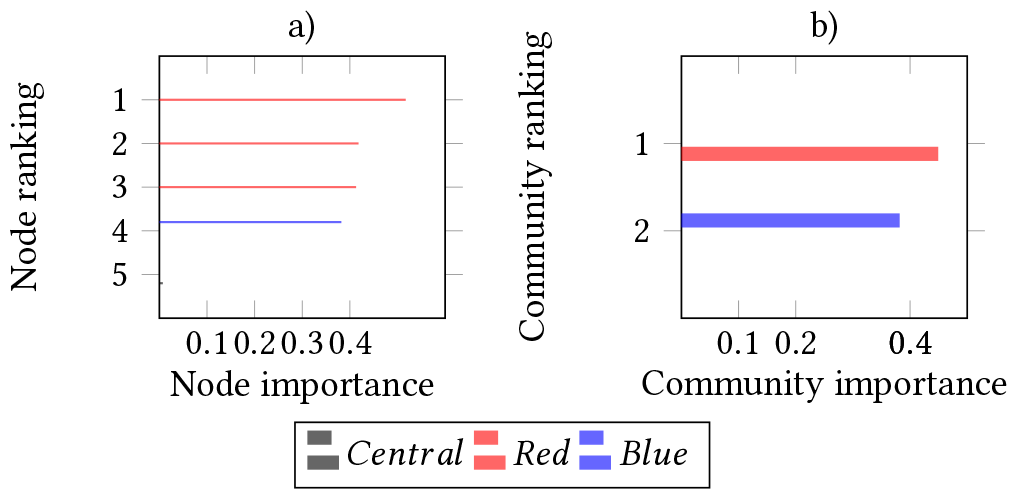
a) Node scores for the central node in the graph in Fig. 7b), when using a GNN to predict node-wise labels. Node bars receive the color of the community to which each node belongs. The central node is colored in black. b) Aggregated scores for the communities in Fig. 7.
- *Case B*, where the three-node community and the central node are set to 0.9 labels, while the node in the other community has a label with a value of 0.1. In this case, the objective is to observe how the three-node community presents a high influence on the behavior of the central node.

The label values of 0.1 and 0.9 represent low and high probability values that could simulate a node classification problem. The values chosen were not zero or one for all nodes to avoid extreme situations, having closer probabilities to a real-world setup.

A four-hop Graph Convolutional Network (GCN) [17] was trained to predict the given labels for all nodes in the graph, reaching *R*^2^ = 0.99 in a random hold-out test set with 20% of the nodes, in both cases. XP-GNN was then applied to explain the GCN for the central node. To minimize the impact of random initializations of the surrogate model, ten surrogate models trained from different random seeds were averaged inside XP-GNN. We trained each initialization for 50 epochs and 20 graph perturbations per epoch (for more information on how XP-GNN works, see Section 4). In Figs.8-9, we can see the ranking distributions for nodes and communities, respectively.

In case A, the most relevant community according to *XP-GNN* is the one-node community, as in this case, this is the community with the closer label values to the node of interest. In case B, the most influential community is the three-node community, as expected since its labels were specifically chosen to be the same as the label of the central node to check whether *XP-GNN* would retrieve this influence. Consequently, this example shows how *XP-GNN* captures the homophily relations that the communities may exert in the central node of interest. The retrieved configuration values in *Case A* are lower than for *Case B*, as they are related to the total sum of labels of the graph nodes. In *case A*, only two nodes are labeled with value 0.9, while *case B* includes four nodes with that value.

### 5.2 Scalability

The Barabàsi-Albert framework for realistic graph building was applied [33] to test the scalability of XP-GNN. Graphs constructed with the Barabàsi-Albert framework receive as input a node number and an average number of edges between their nodes, with the aim to generate realistic graphs. A set of graphs with that framework was generated with 10 to 50000 nodes and an average of five edges between their nodes. Each node has 16 random features. Five overlapping communities were additionally generated by taking several nodes at random, where there could be nodes assigned to more than one community. We trained GCN models [17] to estimate the degree of each node in the Barabási-Albert graph set. In each graph, we explained with XP-GNN the behavior of GCN models in the node with the highest degree. Ten surrogate models were trained for 50 epochs from different random seeds with Adam optimizer, learning rate 0.01, and 20 perturbations per epoch (see Section 4). The coefficients of these models were then averaged. We registered GPU memory usage and runtime on a 32GB GPU for each graph, as displayed in Fig. 10. GPU memory varies linearly with the number of nodes for linear increases in the size of the perturbation masks in XP-GNN, as the number of regression parameters to be learned increases linearly. At the same time, runtime seems to present a logarithmic pattern against perturbation mask size, so XP-GNN seems to scale better in time than in memory. Even if the graphs have an increasing number of nodes, they contain five large communities with few community-wise perturbations to be explored than if there were more communities. This results in a runtime stabilization, resulting in an increase in runtime for larger graphs, but not a drastical one.

**Figure 10.**
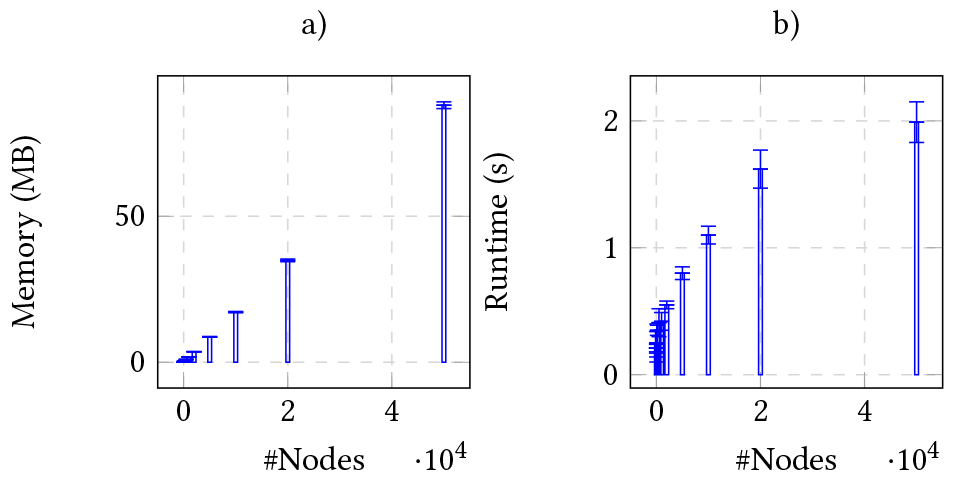
Memory (a) and time (b) requirements for XP-GNN applied on Barabási-Albert graphs, with different number of nodes and constant number of edges per node.

### 5.3 Stability and faithfulness

The model explained by XP-GNN should present a close behavior in a subgraph with only the most relevant communities compared to the input graph, what is known as *faithfulness*. XP-GNN is a *faithful* approach if the predictions of the GNN model for the most relevant communities from XP-GNN approximate the GNN predictions for the same communities in the original complete graph.

To study explanation faithfulness, we sampled a *gene-gene* interaction graph from BIKG [32] (more information on this in Section 6.1). In this graph, nodes represent genes with a series of relations and specific gene features. The genes in the graph group themselves in communities called *pathways* according to their biological function. These gene nodes were labeled as linked to Non-Small Cell Lung Cancer (NSCLC) or not, based on evidence from biomedical databases and previous literature [34–37]. This turned into a node classification problem for a GNN. From the genes in the graph, we explained the EGFR gene node with XP-GNN for an optimized GNN architecture since EGFR presents a strong relation to NSCLC [38]. The biological findings of the XP-GNN explanations of the GNNs applied can be found in Section 6.1. XP-GNN was trained as indicated in that Section, providing a ranking of the least to the most influential nodes and pathways for the behavior of EGFR in NSCLC.

We sequentially added the pathways in the order given in the explanation ranking from XP-GNN, from least to most influential. The EGFR probability for lung cancer was predicted with the GNN in each ablation case, allowing us to observe how the different pathways influence the EGFR predictions. The predicted probability for these subgraphs should converge to the GNN behavior for EGFR in the complete graph. Results are reported in Fig. 11a), including also in 11b) the approximated configuration values for each pathway that is included in each step.

**Figure 11.**
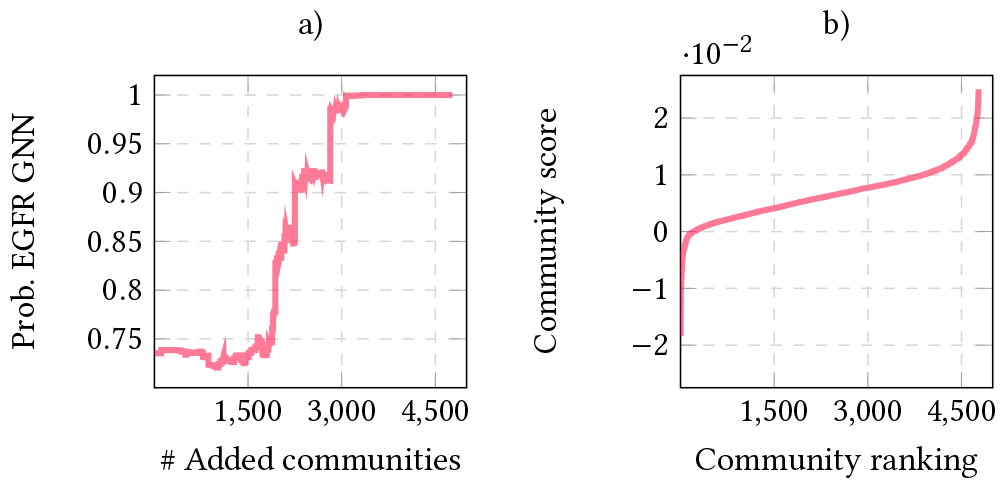
a) Predicted probability of EGFR for NSCLC when including the communities as ranked by XP-GNN. b) XP-GNN community scores in ascending order.

To the left of Fig. 11a) we can observe that graphs with no community information, and only with the isolated node features already reach a probability for EGFR in lung cancer of 0.73, so the node features alone are highly informative. The predictions for EGFR barely change when including the bottom communities of the ranking, as these communities do not present a large influence. Bottom communities also present a configuration value lower than 0.005 (see Fig. 11b)). With the following 1500 communities, the EGFR probability steadily increases, as these communities start to have more influence on the behavior of the GNN in EGFR. The last 1800 communities are the most relevant ones, with a probability of EGFR being linked to NSCLC close to one. We can see that the top 300 communities concentrate the top scores in the upper tail of the curve in Fig. 11b).

In this case, we can state that XP-GNN guarantees faithfulness, as the probabilities for EGFR to be involved in lung cancer increase as the most influential communities for this gene are included (as we move towards the right of Fig. 11a)).

### 5.4 Community size bias

Another aspect analyzed was the potential bias of XP-GNN toward large graph communities. Since large graph communities contain many nodes or edges, it may be natural to think that XP-GNN might place large communities at the top of the explanation rankings, as these could have the potential to modify much more the decisions of a GNN in a node, edge, or graph of interest. To determine whether XP-GNN is biased towards large graph communities, we decided to examine the sizes of the communities in the explanation ranking from the faithfulness experiment in the previous section. Fig. 12 represents the variation in the logged explanation scores for the different communities in that experiment against the respective sizes of these communities. A regression line was fit in the scatter points to find a linear relationship between the log community scores and the community sizes and to determine how strong the potential community size bias could be by studying the slope of the fitting line.

**Figure 12.**
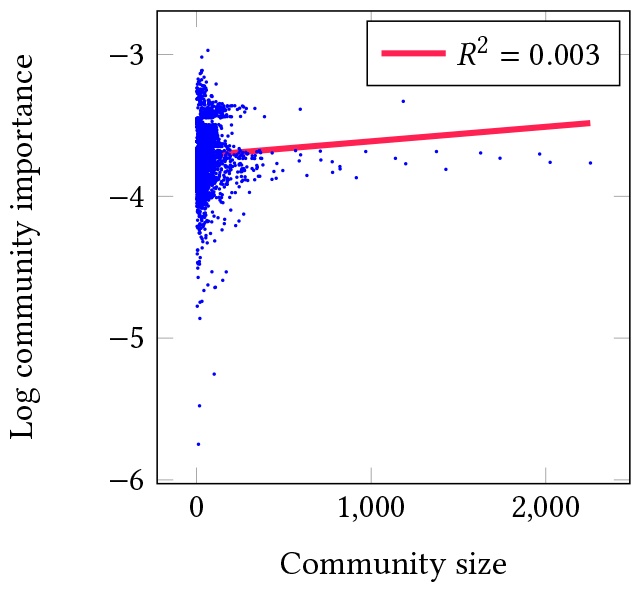
Community size with respect to community importance in XP-GNN, for the same use case as Fig. 11. The plot includes a fitting regression line in red.

The regression line in Fig. 12 presents a *R*^2^ of 0.003, so no linear patterns seem to exist between the logged explanation scores and the community sizes. Furthermore, the line looks almost horizontal, coming closer to a set of communities with a log community importance between -3.5 and -4. Consequently, there does not seem to be a clear bias in XP-GNN where small communities receive a low explanation score and large communities receive a high explanation score, as in that case, the fitting line would be steeper (even if the bias relation would not be linear). Actually, large communities seem to receive intermediate explanation scores, not too small and not too high, given the almost horizontal cluster of points around log community importance values of -3.5. These data points drive most of the line fit, as previously described.

The lack of a clear bias towards large communities at the top of the explanation rankings in XP-GNN is expected. XP-GNN first computes node-wise or edge-wise explanation scores given to the communities in a graph, and the explanation scores are aggregated into community scores through aggregation operators independent of community sizes, such as the mean or the median.

## 6 DRUG DISCOVERY CASE STUDIES

XP-GNN has been evaluated on two use cases in the pharmaceutical industry.

### 6.1 Gene - Disease Prioritization

This use case has been referenced in Section 5.3 to discuss the faith-fulness of XP-GNN. Here we focus more on the findings obtained from the XP-GNN explanations in the same experiment.

#### 6.1.1 Dataset

From the in-house Knowledge Graph BIKG we have sampled a smaller Knowledge Graph with only gene nodes, also known as *gene-gene* graph[32], with 18628 gene nodes, 5712220 gene-gene relation edges, and 4778 gene communities known as *pathways*. The graph edges included gene-gene relations with examples such as *gene-interacts with-gene, gene-modifies-gene*, and *gene-regulates-gene*. Gene nodes contain features retrieved by the Centre of Genomic Research [32]. These genes are arranged into *pathways*, which are gene communities with a common biological function. These *gene pathway communities* are already known, and are not generated by any graph clustering technique, so that the mined communities by *XP-GNN* have a biological meaning. Moreover, gene nodes were labeled according to previous literature as to whether they were involved or not in Non-Small Cell Lung Cancer (NSCLC), resulting in 48 positive gene nodes for lung cancer.

#### 6.1.2 Model

The probability that a gene node would be involved in NSCLC was estimated with a GNN, given the features of each gene and the relations to similar genes. The GNN learned from the existing NSCLC labels in a supervised fashion. A balanced training set was constructed by selecting a random half of the NSCLC-related gene nodes and the same number of random non-NSCLC gene nodes. The test set contained the remaining gene nodes, where we aimed to maximize its ROC AUC with a diverse compendium of GNNs.

The optimal GNN architecture found contained two APPNP layers [20], which combine message-passing techniques with gene node prediction propagation based on *Page Rank* [21], an algorithm that determines the importance of each node based on its connections. Both APPNP layers contained ten propagation iterations with a 0.05 chance for propagation restart from the node of interest. The resulting node embeddings were mapped into the NSCLC probabilities by a two-layer neural network with 84 and 16 layers, respectively. The final metrics provided were a test set ROC AUC of 0.88 and an accuracy of 0.98.

#### 6.1.3 Experimental setup and findings

The gene for the Epidermal Growth Factor Receptor protein (EGFR) is a gene that presents mutations in a significant ratio of NSCLC cases [38]. Consequently, it becomes crucial to determine the most relevant gene nodes and pathways driving its association to NSCLC, applying XP-GNN to the optimal GNN. The XP-GNN explanation scores were averaged between ten different executions to minimize the impact of the random initialization of the linear regression weights (see Section 4). Each initialization was trained for 50 epochs, with 20 graph perturbations per epoch and a learning rate of 0.01. Table 2 contains the top-five communities and nodes and their respective explanation scores.

To determine quantitatively the relevance of the given explanations, as we lack any ground-truth explanation to why a certain gene induces NSCLC, we argue that if EGFR is an NSCLC-related gene node, the top genes and pathway communities in the XP-GNN rankings should also be somehow related to NSCLC. Consequently, we measured the proportion of genes and pathways linked to NSCLC among the top elements in the explanations. In this case, XP-GNN retrieves 47% of the communities in the top 100 as related to lung cancer and 8% of the genes. As a baseline, the total percentage of lung cancer-related communities and genes in the complete ranking is 26% and 0.25%, respectively.

Most of the pathway communities and genes in Table 2 are related in some way or other to EGFR and lung cancer, showing the high degree of relevance of the explanations returned by XP-GNN. EGFR, also known as ERBB1, belongs to the same family as ERBB2, represented in the community *ERBB2 activates PTK6 signaling*. If mutated, ERBB2 might also lead to NSCLC, and the activation of PTK6 (*Protein Tyrosine Kinase 6*) might interfere with the normal function of EGFR, as well [39]. EGFR is also closely related to the FGFR family (*Fibroblast Growth Factor Receptor*), and mutations in members such as FGFR2 and FGRF4 have been frequent in many NSCLC cases [40]. The introduction of community information in the GNN explanation process with XP-GNN seems to detect relevant pathways as these ones. Nevertheless, some unrelated pathways (according to state-of-the-art biological knowledge) were also included among the top 5, according to the current biological knowledge. The presence of unrelated pathways at the top does not necessarily imply that XP-GNN does not mine relevant information, but it could extract information that is still unknown to the current Biology standards.

Regarding the top-5 genes retrieved by XP-GNN, there is also a high degree of relevance, as all genes present some relation with EGFR and NSCLC. SRC was previously proposed as a marker for EGFR-mutated NSCLC, being a proto-oncogene that can drive resistance to current oncology drugs[32]. EREG may participate in *oncogenesis*, contributing to tumor growth and progression [41]. RASGRF1 activates Ras proteins, which also interact with EGFR, being a responder gene to a cancer drug called *Sutinib*[42]. ITPR2 regulates calcium signaling and might interfere with EGFR, too [43]. At last, NRG3 may contribute to tumor growth and progression through its interaction with EGFR [44].

#### 6.1.4 Benchmark for other explanation algorithms

We benchmarked XP-GNN in this setup against the alternative techniques in Table 1. We applied the implementations of these techniques in *GraphXAI*, an open-source library with graph explanation methods [45]. Explanation relevance was measured similarly to the previous section, as the ratio of top-100 gene nodes or pathways in the explanations related to NSCLC. The proportion of NSCLC genes and pathway communities in the full explanation rankings were utilized as *Expected* or baseline measure in the comparison. Since most methods only provide node scores, the pathway community scores were obtained by aggregating the scores for those nodes in the same pathway. Results are displayed in Fig. 13.

**Table 1:**
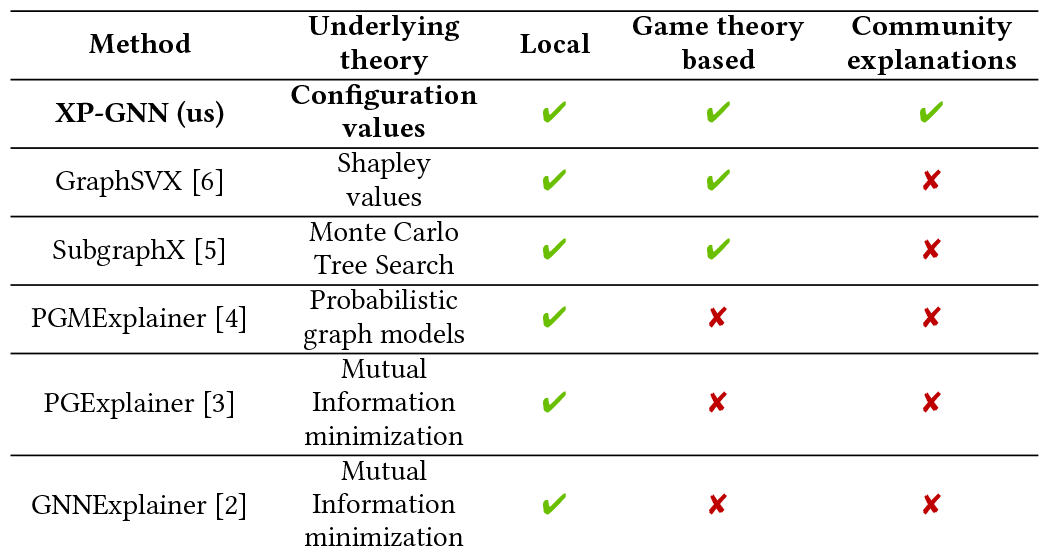
Comparison of XP-GNN against existing GNN explanation techniques.

**Table 2:** The most important communities and nodes that drive EGFR association to lung cancer, according to XP-GNN, in an optimal setup.

| Community | Community score | Gene | Gene score |
| --- | --- | --- | --- |
| Signaling by FGFR2 amplification mutants | 0.0328 | EREG | 0.0469 |
| Maturation of protein E | 0.0304 | ITPR2 | 0.0460 |
| ERBB2 activates PTK6 signaling | 0.0261 | SRC | 0.0455 |
| FGFR4 mutant receptor activation | 0.0260 | RASGRF1 | 0.0403 |
| vRNA synthesis | 0.0254 | NRG3 | 0.0399 |

**Figure 13.**
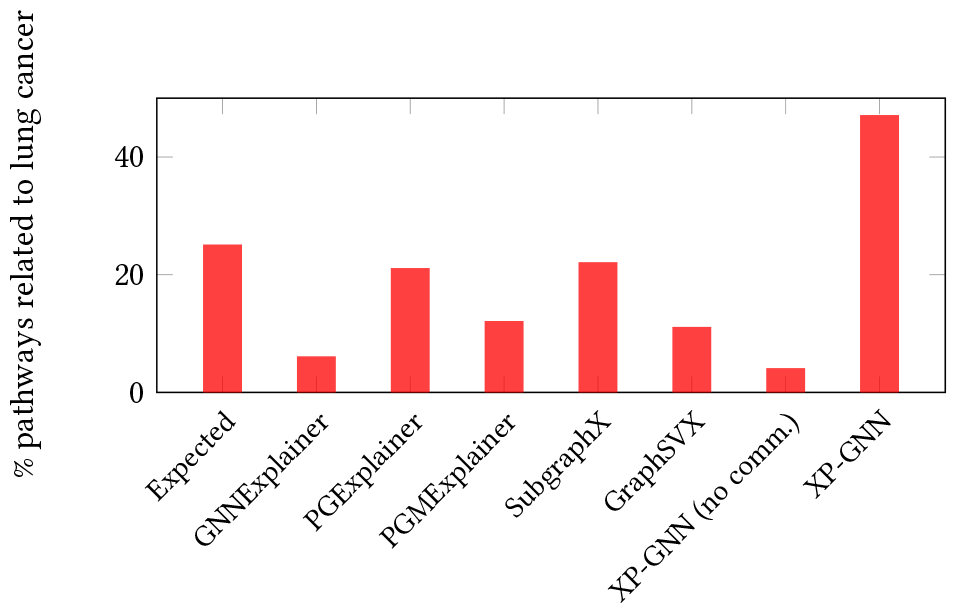
% of lung cancer related pathways for EGFR among the top-100 for each algorithm. The expected % of lung cancer pathways is included as a reference (leftmost bar).

GNNExplainer was executed for 100 epochs and a learning rate of 0.01, while PGExplainer was run for ten epochs and a learning rate of 0.01 in GraphXAI. PGMExplainer was only executed for the chi-square statistics step, without any Probabilistic Graphical Models involved, due to scalability limitations of this technique. SubgraphX was executed with only one MCTS step for a maximum graph size of 1000 neighboring nodes, as it also presented scalability problems. GraphSVX was run from its original implementation, with 1000 graph perturbations and a minimal coalition size of three nodes. Our method, XP-GNN, was additionally executed without prior community awareness, sampling random perturbations without any community criterion (*XP-GNN (no comm*.) in Fig. 13). Among all algorithms, the technique retrieving the most relevant pathways and genes is XP-GNN, as displayed in Fig. 13. In there, most benchmark techniques perform worse than the expected baseline. XP-GNN does not perform so well without prior community information, as *XP-GNN (no comm*.), providing explanations with very low relevance. This result highlights the importance of *a priori* community information in GNN explanations.

### 6.2 Survival prediction

This use case focuses on the estimation of patient survival to NSCLC with a GNN, taking as input the genes that are mutated in each patient and the relations between these genes. For simplicity, the model operates solely with genetic information, but it must be acknowledged that patient age and other environmental factors may play a detrimental role in patient survival. XP-GNN could provide explanations for individual patient survival, with the possibility to be applied in personalized medicine.

#### 6.2.1 Dataset

The *Memorial Sloan Kettering Cancer Center* (*MSKCC*) dataset [46] contains data on mutated genes in 344 NSCLC patients. A *gene-patient* graph was constructed then, linking in both directions each patient node to the gene nodes where it presented mutations (*gene-mutated inpatient, patient-presents mutation ingene*). The genes were also linked between themselves using the same linking edges as in Section 6.1 (*gene-interacts with-gene, gene-modifies-gene*, and *gene-regulates-gene*). The resulting graph consisted of a gene-patient Knowledge Graph with 18628 gene nodes, 344 patient nodes, 5712220 gene-gene edges, and 3683 gene-patient nodes. The graph contains the same 4778 gene pathway communities as in previous Section, using the same gene pathways as in Section 6.1, with an additional community containing all patient nodes.

Each patient was labeled with the follow-up number of months until death or the end of the study. For simplicity, this number was thresholded in six months so that patient nodes were labeled as alive or dead after this period, converting the problem into a node classification task. This criterion resulted in 30.5% of alive patient nodes.

#### 6.2.2 Model

We estimated survival probability on the patient nodes with a wide range of GNN architectures. A random test set was built with 25% of the patient nodes, with a size of 86 patients, where 15 of those patients did survive. ROC AUC and accuracy were estimated as test metrics.

The optimal architecture maximized ROC AUC in the test set, relying on *Relational-Graph Attention modules* (*R-GAT*) [47], where message-passing information between nodes is weighted by attention coefficients while considering different node and edge types. This model reached a ROC AUC of 0.70 and an accuracy of 0.71 in the test set, applying an R-GAT layer with size 16, and a linear layer with size 16. This architecture was trained for 500 epochs and 0.4 dropout probability with a learning rate of 0.1 and Adam optimizer.

#### 6.2.3 Experimental setup and findings

We decided to explain the patient node (*P-0000082-T01-IM3*) since the node presented long survivability with many gene mutations so that XP-GNN could provide more information on the genetic causes of long survivability from the model described above. Table 3 shows the most relevant communities and genes involved in the survivability of this patient.

**Table 3:**
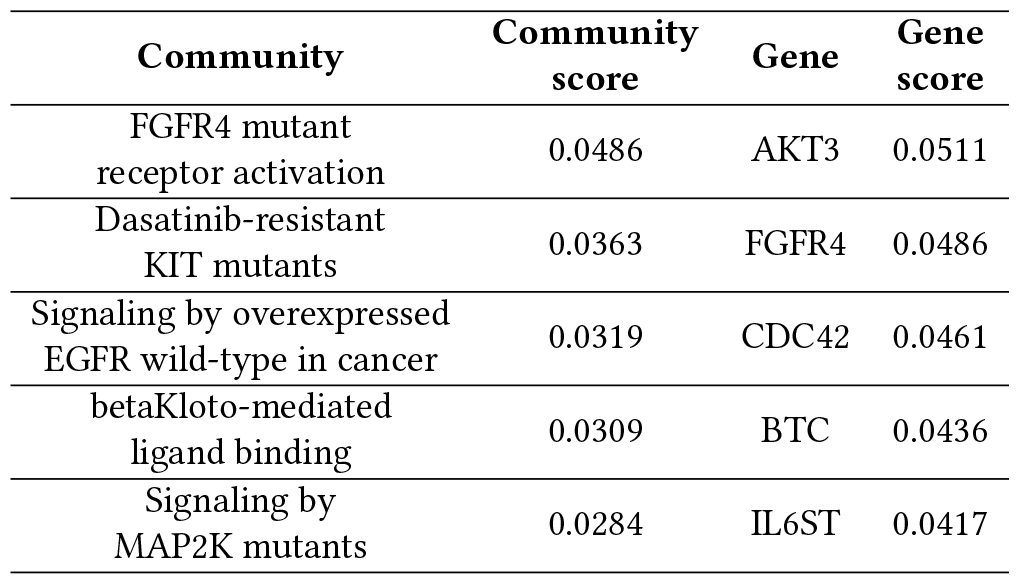
The most important communities and nodes that influence the survival of patient *P-0000082-T01-IM3* in the MSKCC dataset, according to XP-GNN, in the architecture that finds the most related lung cancer information (R-GAT)

As in Section 6.1, the relevance of the top communities and genes in Table 3 was also assessed. In this case, the focus was on communities and genes that could explain the survival to NSCLC of the patient *P-0000082-T01-IM3*, despite its mutations. Thus, it was important to check whether the outputted communities and genes were related to NSCLC, as these could be important factors in cancer survival. Again, *FGFR4 mutant receptor activation* is present in the list, and we already commented that FGFR is a family of genes strongly related to EGFR that might be involved in NSCLC cases. Drug resistance is also a factor that could influence patient survival, in the pathway *Dasatinib-resistant KIT mutants*, although *Dasatinib* is used to treat leukemia rather than NSCLC [48]. EGFR-mediated cancer also plays a role in this explanation, as has already been highlighted in this work, being a target for many drugs against NSCLC [32]. The pathway *betaKlotho-mediated ligand binding* works with Fibroblast Growth Factors, which are closely related to Fibroblast Growth Factor Receptors or FGFRs, the same family of genes present in the pathway *FGFR4 mutant receptor activation* and that could be also involved in NSCLC[49]. Lastly, mutations of MAPK are present in many NSCLC cases, being also quite relevant for the survival of cancer patients[50]. All these communities highlighted by XP-GNN could help explain which pathways could be defective in a specific patient, opening the door to personalized medicine therapies.

The top genes that could be involved in the disease of *P-0000082-T01-IM3* patient are mostly highly relevant and tightly related to NSCLC. For example, FGFR4 appears as well in this last as an individual gene, which should not be surprising, as it is directly or indirectly involved with two of the most important pathways from XP-GNN (*FGFR4 mutated receptor activation* and *betaKlotho-mediated ligand binding*, which we already mentioned that present a relation to NSCLC). This demonstrates that the pathway and gene explanation rankings are highly related. Furthermore, FGFR4 is amplified in NSCLC, being a target for drugs against cholangiocarcinoma and bladder carcinoma [51]. Combinatorial strategies to tackle EGFR-mutated tumors with high FGFR4 activity may be thus beneficial. AKT3 has been shown to influence tumor growth and metastasis in NSCLC [52], being targeted by the small molecules (*ipatasertib, capivasertib*) in late-phase development [53]. CDC42 is a gene involved in cell division, a crucial process for any cancer type [54]. BTC is related to EGFR and lung cancer in a set of cases [55]. Treatments targeting IL6ST may also be a potential therapy for NSCLC, especially for patients treated with immune checkpoint inhibitors [56]. Consequently, the exploitation of community information has also helped in this case to identify lung cancer-related genes and pathways that could be key in determining the survival of the patient of interest.

## 7 CONCLUSIONS AND FUTURE DIRECTIONS

This work presents *XP-GNN*, a novel explanation technique for GNNs that considers the potential overlapping communities of nodes or edges in graphs such as KGs. It is critical to understand the decisions taken by GNNs on top of KGs, especially when a therapeutic choice for a patient is at play. Since the biological entities in the nodes of a KG usually conduct shared functions in graph communities, we consider that explanations based on graph communities may be more realistic than just an isolated set of graph nodes or edges, as many previous explanation techniques address. Since graphs can reach large sizes, it is crucial to have scalable techniques for the viability of graph explanations. With this in mind, XP-GNN approximates the node-wise or edge-wise configuration values [29] of a KG analyzed with a GNN while trying to allow for scalability. XP-GNN completes this by perturbing the input graph into many parallel perturbations based on the present graph communities. The explanations are derived after fitting a weighted regression surrogate model for the GNN outputs of the perturbed graphs [28].

For simplicity, XP-GNN is tested first in a toy example. In there, XP-GNN returns the expected explanations for the basic communities set up, assuming homophily between neighboring nodes. Regarding scalability with Barabàsi-Albert graphs of increasing size, XP-GNN shows a linear pattern for GPU memory, and a logarithmic trend for runtime, showing a promising potential to scale in runtime for large applications. XP-GNN proves to be faithful in a gene classification problem for lung cancer, as the communities at the top of the explanation ranking for a gene of interest can alone provide closer predictions for that gene compared to the complete graph. In the same problem, XP-GNN does not seem to provide high explanation scores for communities with larger sizes, so XP-GNN seems to be an unbiased technique for community size.

For two real-world problems in the pharmaceutical industry, XP-GNN also succeeds in returning relevant nodes (*genes*) and communities (*pathways*) of interest. We defined the relevance of the XP-GNN explanations with the following rationale: if the graph element that was explained was related to a disease of interest, its explanations should also be related to that disease. Consequently, explanation relevance was measured as the ratio of nodes and communities involved with that disease among the top-100 list of ranked genes and communities. The ratio was compared to the total fraction of genes and graph communities that were related to the disease of interest in the complete ranking. XP-GNN provided higher ratios of relevant genes and pathways in the top 100 than in the complete ranking. When XP-GNN is compared to other benchmark techniques, it provided more relevant explanations, too. Domain experts also certified that many of the genes and pathways at the top of the explanation rankings were involved in the disease of interest.

XP-GNN also presents some limitations to be addressed in future work:

- The quantitative metrics used for explanation relevance in XP-GNN are arbitrary, as no exact ground-truth explanation is known. Alternative metrics to the ratio of disease-related pathways and genes in the top of the explanation rankings could have been applied.
- XP-GNN presents some randomness in the outputted explanations as the input graph may be perturbed in different communities in each execution. Moreover, the initial parameters of the surrogate model are randomized. Consequently, it is recommended to average the outputs from a few XP-GNN initializations to reach more stable explanations.
- XP-GNN does not seem to perform well without prior community information. Thus, we highly recommend applying XP-GNN when a community structure is known.
- Even if XP-GNN scales well on runtime, it still presents issues for GPU memory. One may require GPUs with dozens of GB to process KGs with thousands of nodes and millions of edges.
- It is unknown how XP-GNN may scale for KGs with many node types and edge types. Due to internal company reasons, we had to prioritize the application of XP-GNN on large KGs but with few types of nodes and edges.

Future efforts may focus on improved scalability and enhanced abilities of XP-GNN. Communities in KGs can be hierarchically arranged, with some KGs containing *communities of communities*. We may extract more realistic explanations by modeling these hierarchies with more advanced cooperative game theory tools, such as the *Winter values*[57]. Another issue to tackle is explanation *redundancy*, where the explanation rankings may contain communities with similar biological functions at the top, thus limiting explanation diversity. There is some previous work on redundancy filtering we could base for XP-GNN as well [58].

## Notes

https://github.com/andres2631996/bikg_graph_explainability_public

### Competing Interest Statement

A.M.M and D.P were employees and shareholders of AstraZeneca in the past. S.N. and M.U. are currently employed by AstraZeneca, and also hold shares.

