## Supplementary material for "Community-aware explanations in knowledge graphs with XP-GNN"

### A SUPPLEMENTARY INFORMATION

Fig.14 depicts XP-GNN explanation relevance for EGFR related to Non-Small Cell Lung Cancer (NSCLC). Explanations cover different architectures, following the setup introduced in Section 6.1. We can observe that GIN[19] and APPNP[20] provide the most relevant explanations, which also coincides with the top-performing architectures, as introduced in Table 4. GAT[18] and R-GAT[47] also provide relevant explanations in the gene rankings. It seems that the better the architecture performs in the test set, the more relevant explanations are retrieved, which makes sense, as the GNN captures the *real* reasons for classifying EGFR as a lung cancer-related gene.

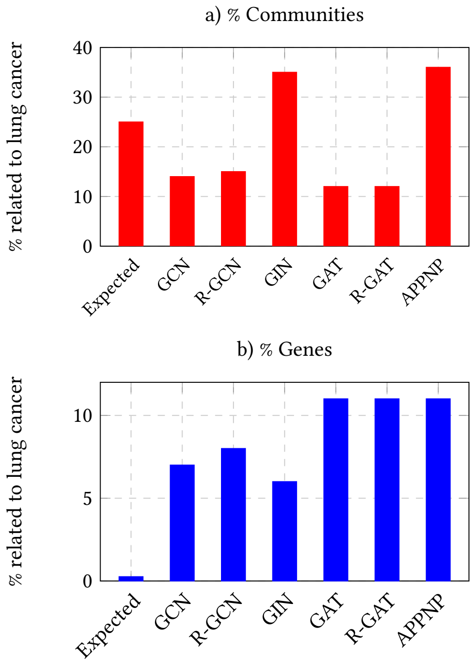


**Figure 14: % of lung cancer-related communities in a) and genes in b) for EGFR among the top 100 for each architecture. We include the expected % of lung cancer communities and genes in the overall rankings as a reference**

Fig.13 represents gene explanation relevance for the same setup as Fig.14, for different graph explanation techniques using an optimal APPNP architecture[20]. As in the community relevance setup in Fig.13, XP-GNN is the method providing the most relevant gene explanations. All techniques provide more positive genes in the top 100 than the overall positive ratio in the explanation rankings, being relevant up to some point. GNNExplainer[2] and SubgraphX[5] seem to be the best-performing techniques from previous literature, but still far from our approach, which shows that the introduction of prior community information helps in providing more reliable and realistic information. This is emphasized by the fact that XP-GNN performance when not inputting any community information at all (in *XP-GNN (no comm.)*) drops drastically. However, we should acknowledge that some methods are limited in terms of scalability. Techniques such as PGMExplainer[4] and SubgraphX[5] did not perform optimally because the gene-gene graph was large to afford a proper execution. Nevertheless, these methods may provide more relevant explanations in simpler graphs.

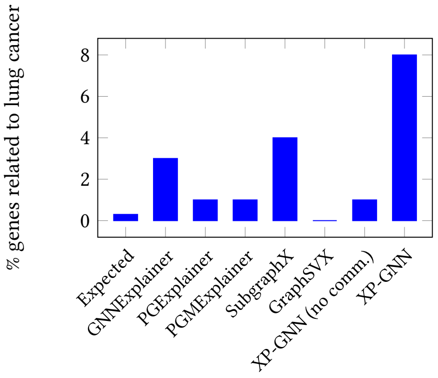


**Figure 15: % of lung cancer related genes for EGFR among the top-100 for each algorithm. We include the expected % of lung cancer pathways as a reference**

Fig.15 shows explanation relevance with XP-GNN for patient *P-0000082-T01-IM3* surviving Non-Small Cell Lung Cancer (NSCLC), for different architectures. The setup used is described in Section 6.2. We quantify explanation relevance as the ratio of communities and genes associated with lung cancer within the top 100 from each explanation ranking. In this case, the best-performing architectures are GAT[18] and R-GAT[47]. R-GAT performs the best in the test set, according to Table 6, confirming our hypothesis that better-performing models get more relevant explanations as they learn from more realistic data.

Table 4 shows the most relevant communities and genes for different architectures, including their relevance as lung cancer-related information ratio in the top 100 of the rankings, and performance in the test set, for the setup in Section 6.1. The explanations between related models, as GCN[17] and R-GCN[59] or GAT[18] and R-GAT[47], are pretty similar, denoting that similar mechanisms of information processing in the GNN receive similar explanations. All explanations seem to contain lung cancer-related concepts, although only GIN[19] and APPNP[20] seem to provide many lung cancer-related concepts.

Table 5 presents the most relevant communities and genes for benchmark methods, including XP-GNN lacking community information. The architecture applied is an APPNP[20], as described in Section 6.1. It also presents explanation relevance. XP-GNN seems to provide more lung cancer-related concepts than previous methods, especially compared to XP-GNN without community information. Previous techniques are influenced by poor performance in large graphs, but they could work better in smaller graphs. This is a question to leave for future research.

Table 6 reflects the test performance scores, relevance scores, and more relevant communities and genes for patient *P-0000082-T01-IM3*, with different architectures. The setup followed is described in Section 6.2. As stated before, the best-performing architectures,

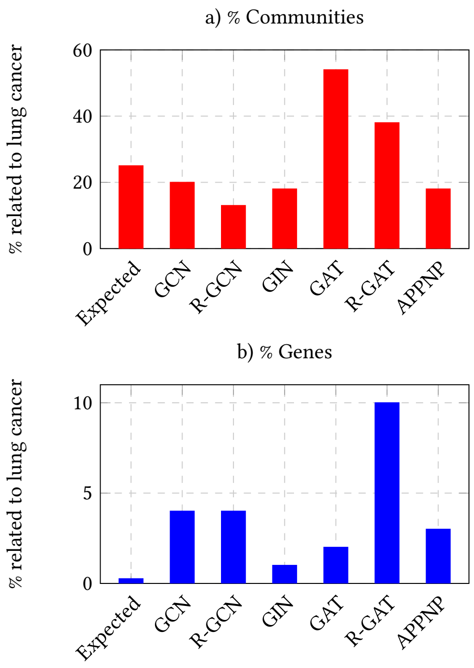


**Figure 16: % of lung cancer-related communities in a) and genes in b) for the patient *P-0000082-T01-IM3* in MSKCC dataset, in the top-100, for several GNN architectures. We include the expected % of lung cancer communities and genes in the overall rankings as a reference**

GAT[18] and R-GAT[47] seem to be the ones with more cancer-related concepts for the survived patient. Additionally, similar architectures, as GCN[17] and R-GCN[59] or GAT[18] and R-GAT[47] present similar explanations between them, as expected.

**Table 4: The most relevant communities that influence EGFR to be associated to lung cancer, for several GNN architectures in XP-GNN. Results are averaged among three random seeds**

| Method | GCN [17] | R-GCN [59] | GIN [19] | GAT [18] | R-GAT [47] | APPNP [20] |
|---|---|---|---|---|---|---|
| ROC AUC | 0.91 | 0.87 | 0.92 | 0.85 | 0.87 | 0.88 |
| Accuracy | 0.71 | 0.68 | 0.76 | 0.75 | 0.69 | 0.98 |
| Comm. 1 | Maturation of protein E | Maturation of protein E | Synthesis and processing of ENV and VPU | MPS IIID Sanfilippo syndrome D | MPS IIID Sanfilippo syndrome D | Dasatinib-resistant KIT mutants |
| Comm. 2 | Synthesis and processing of ENV and VPU | Synthesis and processing of ENV and VPU | Defective SLC2A2 causes Fanconi-Bickel syndrome | Maturation of protein E | Maturation of protein E | Signaling by FGFR1 amplification mutants |
| Comm. 3 | Activation of AP-1 family of transcription factors | SCF FBW7 complex | Maturation of protein E | Butyrate-induced histone acetylation | Phospho-PLA2 pathway | FGFR4 mutant receptor activation |
| Comm. 4 | SCF FBW7 complex | Activation of AP-1 family of transcription factors | SCF FBW7 complex | Phospho-PLA2 pathway | Butyrate-induced histone acetylation | Signaling by FGFR2 amplification mutants |
| Comm. 5 | Phospho-PLA2 pathway | Loss of function of FBWX7 in cancer and NOTCH1 signaling | Loss of function of FBWX7 in cancer and NOTCH1 signaling | AKT-mediated inactivation of FOXO1A | Signaling by overexpressed wild-type EGFR in cancer | GRB7 events in ERBB2 signaling |
| Gene 1 | CSNK2A1 | NFKB1 | CSNK2A1 | MYC | MYC | ERBB2 |
| Gene 2 | CREBBP | CSNK2A1 | HDAC3 | AKT1 | SRC | PSMB7 |
| Gene 3 | NFKB1 | MYL12A | NFKB1 | SRC | RAC1 | SRC |
| Gene 4 | MAPK10 | MAPK10 | MYL12A | NFKB1 | NFKB1 | YWHAZ |
| Gene 5 | MYL12A | CREBBP | PPP2R1A | RAC1 | AKT1 | NRAS |
| % related comms. in top 100 | 14 | 15 | 35 | 12 | 12 | 36 |
| % related genes in top 100 | 7 | 8 | 6 | 11 | 11 | 11 |

**Table 5: The most important pathways that influence that EGFR as associated to lung cancer, for several explanation methods. Results are averaged among ten random seeds**

| **Method** | **GNNExplainer [2]** | **PGExplainer [3]** | **PGMExplainer [4]** | **SubgraphX [5]** | **GraphSVX [6]** | **XP-GNN (no communities)** | **XP-GNN** |
|---|---|---|---|---|---|---|---|
| **Comm. 1** | Butyrate-induced histone acetylation | Defective translocation of RB1 mutants to the nucleus | Signaling by overexpressed wild-type EGFR in cancer | Signaling by MAP2K mutants | Phosphatidyl-coline (PC) biosynthesis, PE => PC | Defective MAN1B1 causes MRT15 | Signaling by FGFR2 amplification mutants |
| **Comm. 2** | Defective CYP27B1 causes VDDR1A | FA core complex | RPIA deficiency: failed conversion of R5P to RU5P | MEK activation | Defective SLC12A3 causes Gitelman syndrome (GS) | Biosynthesis of DPAn-3 derived protectins and resolvins | Maturation of protein E |
| **Comm. 3** | Defective FMO3 causes trimethylaminuria (TMAU) | ATM mediated phosphorylation of repair proteins | Defective ACY1 causes encefalopathy | Evasion of oncogene induced senescence due to p14ARF defects | Defective SLC17A5 causes Salla disease (SD) and ISSD | Metabolism of ingested MeSeO2H into MeSeH | ERBB2 activates PTK6 signaling |
| **Comm. 4** | Defective UGT1A1 causes hyperbilirubinemia | vRNA synthesis | Biosynthesis of DPAn-3-derived 13-series resolvins | PTEN loss of function in cancer | Variant SLC6A14 may confer susceptibility towards obesity | Defective HK1 causes hexokinase (HK) deficiency | FGFR4 mutant receptor activation |
| **Comm. 5** | Defective CYP7B1 causes SPG5A and CBAS3 | Loss of function of TP53 in cancer due to loss of tetramerization ability | Defective MAN1B1 causes MRT15 | RAS GTPase cycle mutants | Defective GSS causes Gluthatione synthetase (GSS) deficiency | Defective SLCO2A1 causes primary autosomal recessive hypertrophic osteoartropathy 2 (PHOAR2) | vRNA synthesis |
| **Gene** | C17orf97 | NFKB1 | EGFR | EGFR | MAP4K3 | BRCA2 | EREG |
| **Gene 2** | SPATA8 | CSNK2A1 | TCIRG1 | SRC | PATL1 | NUP88 | ITPR2 |
| **Gene 3** | KRTAP9-1 | MYL12A | ZNF716 | MAPK7 | TMEM165 | FBXL3 | SRC |
| **Gene 4** | PTER | MAPK10 | PDPR | MAPK13 | GAL3ST2 | RPA3 | RASGRF1 |
| **Gene 5** | WNT11 | CREBBP | FBXL20 | MAPK14 | AMHR2 | FAM76A | NRG3 |
| **% related comms. in top 100** | 6 | 21 | 12 | 12 | 11 | 4 | 47 |
| **% related genes in top 100** | 3 | 1 | 1 | 1 | 0 | 1 | 8 |

**Table 6: The most relevant communities that influence the survival of patient *P-0000082-T01-IM3* in MSKCC dataset, for several GNN architectures. Results are averaged among three random seeds**

| **Method** | **GCN [17]** | **R-GCN [59]** | **GIN [19]** | **GAT [18]** | **R-GAT [47]** | **APPNP [20]** |
|---|---|---|---|---|---|---|
| **ROC AUC** | 0.65 | 0.64 | 0.64 | 0.60 | 0.70 | 0.59 |
| **Accuracy** | 0.69 | 0.70 | 0.56 | 0.57 | 0.71 | 0.71 |
| **Comm. 1** | Loss of function of TP53 in cancer due to loss of tetramerization ability | Loss of function of TP53 in cancer due to loss of tetramerization ability | Evasion of oncogene-induced senescence due to p14ARF defects | Signaling by overexpressed wild-type EGFR in cancer | FGFR4 mutant receptor activation | Influenza virus induced apoptosis |
| **Comm. 2** | AKT-mediated inactivation of FOXO1A | PTEN loss of function in cancer | Signaling to ERK5 | RAS GTPase cycle mutants | Dasatinib-resistant KIT mutants | RNA polymerase II, eukaryotes |
| **Comm. 3** | Activation of AP-1 family of transcription factors | Transcriptional activation of cell cycle inhibitor p21 | E-cadherin signaling events | betaKlotho-mediated ligand binding | Signaling by overexpressed wild-type EGFR in cancer | Loss of function of TP53 in cancer due to loss of tetramerization ability |
| **Comm. 4** | RNA polymerase II, eukaryotes | MET activates PTPN11 | Synthesis of PIPs in the nucleus | FGFR4 mutant receptor activation | betaKlotho-mediated ligand binding | Mechanism of pioglitazone and rosiglitazon action in diabetes mellitus type 2 |
| **Comm. 5** | Maturation of protein E | Imatinib-resistant PDGFR mutants | Defective DNA double strand break response due to BARD1 loss of function | MEK activation | Signaling by MAP2K mutants | Defective SLC2A2 causes Fanconi-Bickel syndrome (FBS) |
| **Gene 1** | RELA | KL | RPL18 | PPP2R1A | AKT3 | TGFB1 |
| **Gene 2** | MAPK3 | ADCY1 | RPL36A | PPP2CA | FGFR4 | ACTB |
| **Gene 3** | JUN | CDC42 | RASAL3 | HSP90B1 | CDC42 | RELA |
| **Gene 4** | RPS27A | EGF | RPS23 | PRKCQ | BTC | NFNB1 |
| **Gene 5** | PIK3CB | TGFA | MAPK13 | EGF | IL6ST | TNF |
| **% related comms. in top 100** | 20 | 13 | 18 | 54 | 38 | 18 |
| **% related genes in top 100** | 4 | 4 | 1 | 2 | 10 | 3 |

As part of the Supplementary Information, we also attach the full reports from the most relevant genes in the gene-disease use case in Section 6.1, and in Section 6.2. They were generated with the software *OncoEnrichR*, for gene enrichment analysis [60], from the top genes in each use case. These reports were then used by field experts to evaluate the relevance of the genes provided by XP-GNN in the explanations.
